# Evolution of dosage-sensitive genes by tissue-restricted expression changes

**DOI:** 10.1101/2024.10.17.618887

**Authors:** Alan M. Rice, Yuanshuo Li, Pauric Donnelly, Aoife McLysaght

## Abstract

Dosage-sensitive genes have characteristic patterns of evolution that include being refractory to small-scale duplication, depleted on human benign copy number variants (CNVs) and enriched on pathogenic CNVs. This intolerance to copy number change is likely due to an expression constraint that exists in one or more tissues. While genomic copy number changes alter the encompassed genes’ expression across all tissues, expression quantitative trait loci (eQTLs) –genomic regions harbouring sequence variants that influence the expression level of one or more genes– can act in a tissue-specific manner. In this work we examine expression variation of presumed dosage-sensitive and non-dosage-sensitive genes to discover how the locus duplicability constraints translate into gene expression constraints. Here we test the hypothesis that expression changes due to the presence of eQTLs acting in unconstrained tissues will not be deleterious and thus allow dosage-sensitive genes to vary expression while obeying constraints in other tissues. Using eQTLs across 48 human tissues from The Genotype-Tissue Expression (GTEx) project, we find that dosage-sensitive genes are enriched for being affected by eQTLs and that the eQTLs affecting dosage-sensitive genes are biased towards having narrow tissue-specificity with these genes having fewer eQTL-affected tissues than non-dosage-sensitive genes. Additionally, we find that dosage-sensitive genes are depleted for being affected by broad tissue breadth eQTLs, likely due to the increased chance of these eQTLs conflicting with expression constraints and being removed by purifying selection. These patterns suggest that dosage-sensitivity shapes the evolution of these genes by precluding copy number evolution and restricting their evolutionary trajectories to changes in expression regulation compatible with their functional constraints. Thus deeper interpretation of the patterns of constraints can be informative of the temporal or spatial location of the gene dosage sensitivity and contribute to our understanding of functional genomics.

**Author summary:** Gene duplication is an important and powerful evolutionary force that is responsible for the expansion of the coding capacity of genomes ultimately resulting in great genetic novelty. However, the opportunity for this evolutionary change can be limited by dosage constraints on some genes, meaning they are not normally duplicable, except in a balanced, whole genome event. This results in important, biologically relevant, differences between genes that are retained from whole genome duplication events versus those retained from small scale duplications, especially in terms of dosage sensitivity. We explored how the different dosage sensitivity in these sets of genes relates to quantitative expression variation present in populations. We found that while dosage-sensitive genes are more likely to have their expression levels influenced by genetic variation, these changes are often specific a small number of tissues. In contrast, genes that are less sensitive to dosage changes show greater variation in expression levels across multiple tissues. Our findings suggest that dosage-sensitive genes evolve through fine-tuned adjustments in their expression levels in specific tissues, thus bypassing constraints operating on other tissues. This understanding sheds light on how dosage-sensitive genes evolve and could have implications for understanding human diseases caused by these genes.

## Introduction

Gene duplication is a powerful force that is responsible for a great deal of evolutionary innovation (Prince and Pickett 2002). Evolutionary duplications are broadly classified into those that emerge from whole genome duplication (WGD), with the remainder grouped as small-scale duplications (SSDs). At a population genetics level, duplications are observed as copy number variants (CNVs) that are polymorphic between individuals. While it might be tempting to think that a duplicate is a duplicate, a large and growing body of evidence points to the different properties of genes that are retained in duplicate after WGD (termed ‘ohnologs’) and those that are commonly observed as SSDs, with ohnologs being generally longer, more highly expressed, slower evolving, and more associated with disease (Makino and McLysaght 2010; Vance and McLysaght 2023). Additionally, retained ohnologs and SSDs have clear differences in terms of dosage-sensitivity, which manifests as copy number constraints.

Dosage sensitive genes are an important subset of genes in our genome that include many developmental genes, protein complex members and transcription factors among others (Birchler and Veitia 2012; Maere et al. 2005). They are described for the relationship between gene dosage and functionality, where, broadly speaking, a different dosage will cause a change in functional outcome or even a malfunction (Veitia 2002). In human genetics this is observed as genes with a phenotype (especially a disease phenotype) when the copy number is altered through structural variation (Zhang et al. 2009; Cooper et al. 2011). Over evolutionary timescales this creates obvious constraints. These constraints leave distinctive traces in the evolutionary patterns of dosage sensitive genes – they are observed as genes that are refractory to the otherwise pervasive process of gene duplication (Papp, Pál, and Hurst 2003), except whole genome duplication, following which they are disproportionately retained (Birchler, Riddle, et al. 2005; Makino and McLysaght 2010; Birchler, Bhadra, et al. 2001; Tasdighian et al. 2017; Goût and Lynch 2015).

Dosage sensitivity also shapes the evolutionary trajectory of the respective genes in various other ways. Previous work has explored gene dosage sensitivity in the context of evolutionary duplicability, and population-level copy number variation (Papp, Pál, and Hurst 2003; Makino and McLysaght 2010; Rice and McLysaght 2017; Schuster-Böckler, Conrad, and Bateman 2010; Goût and Lynch 2015; Gout et al. 2010), as well as other forms of functional constraint (Xie et al. 2016).

There are fewer studies that explicitly test expression evolution of dosage sensitive genes, and those that there are, suggest that the constraints observed on genomic and coding sequence features extend to expression features. Genes whose proteins are members of protein complexes are likely to be dosage sensitive (Papp, Pál, and Hurst 2003), and are also less likely to vary in expression between individuals (Schuster-Böckler, Conrad, and Bateman 2010). Furthermore, genes with protein-protein interactions are more constrained in their regulatory evolution and have less expression polymorphism within populations (Lemos, Meiklejohn, and Hartl 2004).

The availability of large expression quantitative trait locus (eQTL; genomic regions harbouring sequence variants that influence the expression level of one or more genes (Albert and Kruglyak 2015)) datasets for humans and many other species, means that it is now possible to test the relationship between dosage constraints and expression evolution constraints in a more comprehensive way and at scale (Morley et al. 2004; Cheung et al. 2005; Stranger, Forrest, et al. 2005; Stranger, Nica, et al. 2007; West et al. 2007; Dimas et al. 2009; Kelly et al. 2012; Massouras et al. 2012; GTEx Consortium 2017).

The Genotype-Tissue Expression (GTEx) project (GTEx Consortium 2017) has characterised eQTLs across a diverse range of human tissues. In Release V7, 95.5% (18,199/19,067) of protein-coding genes tested had their expression influenced by at least one eQTL. Given that such a high proportion of the genome experiences this type of expression variation in control individuals, the majority of the genome must be able to tolerate some amount of mRNA level change without obvious deleterious consequences. However, in combination with genome-wide association studies, eQTLs have been used to elucidate further the pathophysiology of many disease phenotypes. To date eQTLs have been associated with human diseases including asthma, autoimmune disorders, diabetes, numerous cancers, Parkinson’s disease, and other brain disorders (see Table 1 in Albert and Kruglyak 2015). Additionally, eQTLs have been shown to be under increased purifying selection with gene age where young, primate-specific genes are enriched for eQTLs, having higher effect size and influencing expression in more tissues (Popadin et al. 2014). Therefore, the effect of eQTLs on gene expression and association with important traits makes them of great interest, especially in the context of genes with known expression constraints.

**Table 1.**
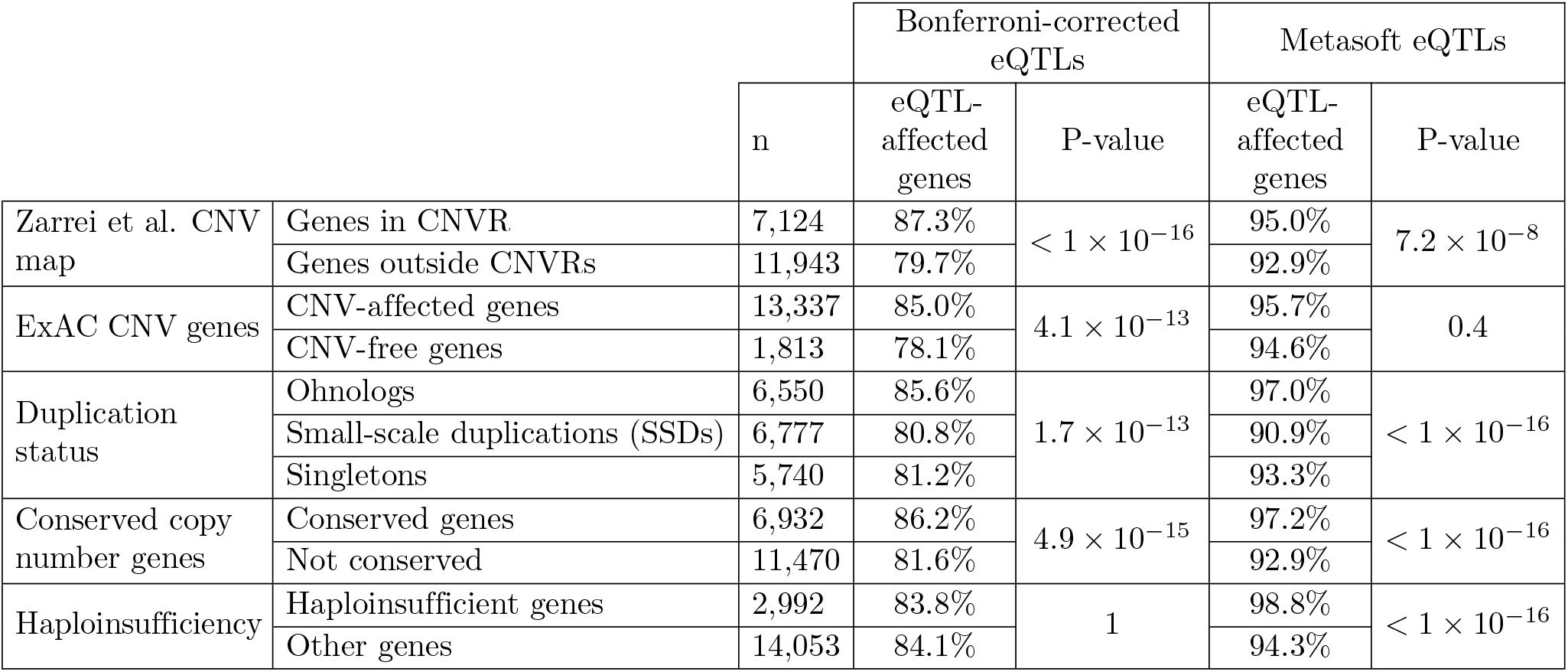
eQTL enrichment of gene groups. P-values for *χ*^2^ tests are Bonferroni-corrected for multiple tests.

Here, we investigated the patterns of eQTLs affecting different types of duplicate genes in the conext of their propensity for dosage-sensitivity. Contrary to the simplistic expectation that ohnologs and other categories of dosage-sensitive genes should be depleted for this variation, we found that these genes are enriched for eQTLs. However, they have fewer eQTL-affected tissues than other genes, as the eQTLs that affect dosage-sensitive genes are more tissue-specific. Dosage-sensitive genes are depleted for broad tissue breadth eQTLs which are likely removed by purifying selection as they conflict with expression constraints. This is consistent with the view that, by contrast to genomic duplications, more subtle dosage changes to dosage sensitive genes may be effectively neutral (Birchler and Veitia 2012). This supports a model where the evolution of dosage sensitive genes is constrained into the comparatively narrow path of tissue-restricted expression changes that do not clash with the essential dosage sensitivity either due to the effect size, or the tissue affected. This opens up the possibility of a deeper understanding of the underlying nature of the dosage sensitivity.

## Results

### Ohnologs are often affected by eQTLs, but they are more distinct between tissues

We gathered two high-confidence sets of eQTLs from the Genotype-Tissue Expression (GTEx) project V7 (GTEx Consortium 2017). One contains significant single tissue SNP-gene associations for 48 tissues corrected for testing across multiple tissues (Supp Figure **??** hereafter ‘Bonferroni-corrected eQTLs’). The other results from a GTEx Consortium meta-analysis using Metasoft which increases eQTL detection power by considering data across tissues together and calculates a posterior probability of an eQTL being present in each tissue (Han and Eskin 2012) (hereafter ‘Metasoft eQTLs’). This latter approach is particularly useful for increasing power in tissues with smaller sample sizes (GTEx Consortium 2017). A comparison of the Bonferroni-corrected eQTL dataset and the Metasoft eQTL dataset can be seen in Supp Figure **??**.

We sought to consider the role of eQTL-based expression variation in the context of gene duplicability and dosage-sensitivity. Assembling a list of dosage sensitive genes is generally based on indirect evidence. Previous work has shown that ohnologs are enriched for dosage-sensitive genes (Makino and McLysaght 2010), as are genes that are conserved in copy number across mammals (Rice and McLysaght 2017), whereas genes that are found as small-scale duplications (SSDs) or present in (benign) CNVs are unlikely to be dosage sensitive (Makino, McLysaght, and Kawata 2013),. Each of these evolutionary genomic metrics is reflecting dosage sensitivity, though perhaps in slightly different ways. There is a good deal of overlap between the various categories (Supp Figure **??**), but they are capturing slightly different information. For example, a given gene may never be observed in a CNV in healthy individuals because it is itself highly dosage sensitive, or because it is closely linked to a dosage-sensitive gene, or because it lies in a region of chromosome less prone to CNV events. This means that while the genes within CNV regions in healthy individuals are unlikely to be dosage-sensitive, the genes outside those regions will be a mix of dosage-sensitive and non-dosage-sensitive genes. Similarly, ohnologs are biased towards dosage sensitive genes, but are neither exclusively nor uniquely dosage sensitive. While noting these caveats, throughout this work we use these sets of genes as proxies for dosage sensitive genes.

Genes that are observed in CNVs in healthy individuals are unlikely to be strongly dosage-sensitive therefore we expect that CNV-affected genes will have little expression constraint. We find support for this simple expectation from examination of genes within CNV regions (CNVRs). Taking recurrent CNVRs described in the inclusive CNV map published by Zarrei et al. (2015), as well as CNV-affected genes across ∼60,000 exomes analysed by the Exome Aggregation Consortium (ExAC) (Ruderfer et al. 2016) we find that genes found within CNVs are enriched for being affected by eQTLs relative to genes outside CNVs (Figure 1A and Table 1). This pattern is consistent for both the Bonferroni-corrected eQTLs and the Metasoft eQTLs but the latter is not significant for the ExAC CNVs (Table 1 and Supp Figure **??**). Genes in CNVs also have a larger absolute number of SNPs and a larger proportion of those that are found as significant eQTLs (see Supplementary Information).

**Figure 1.**
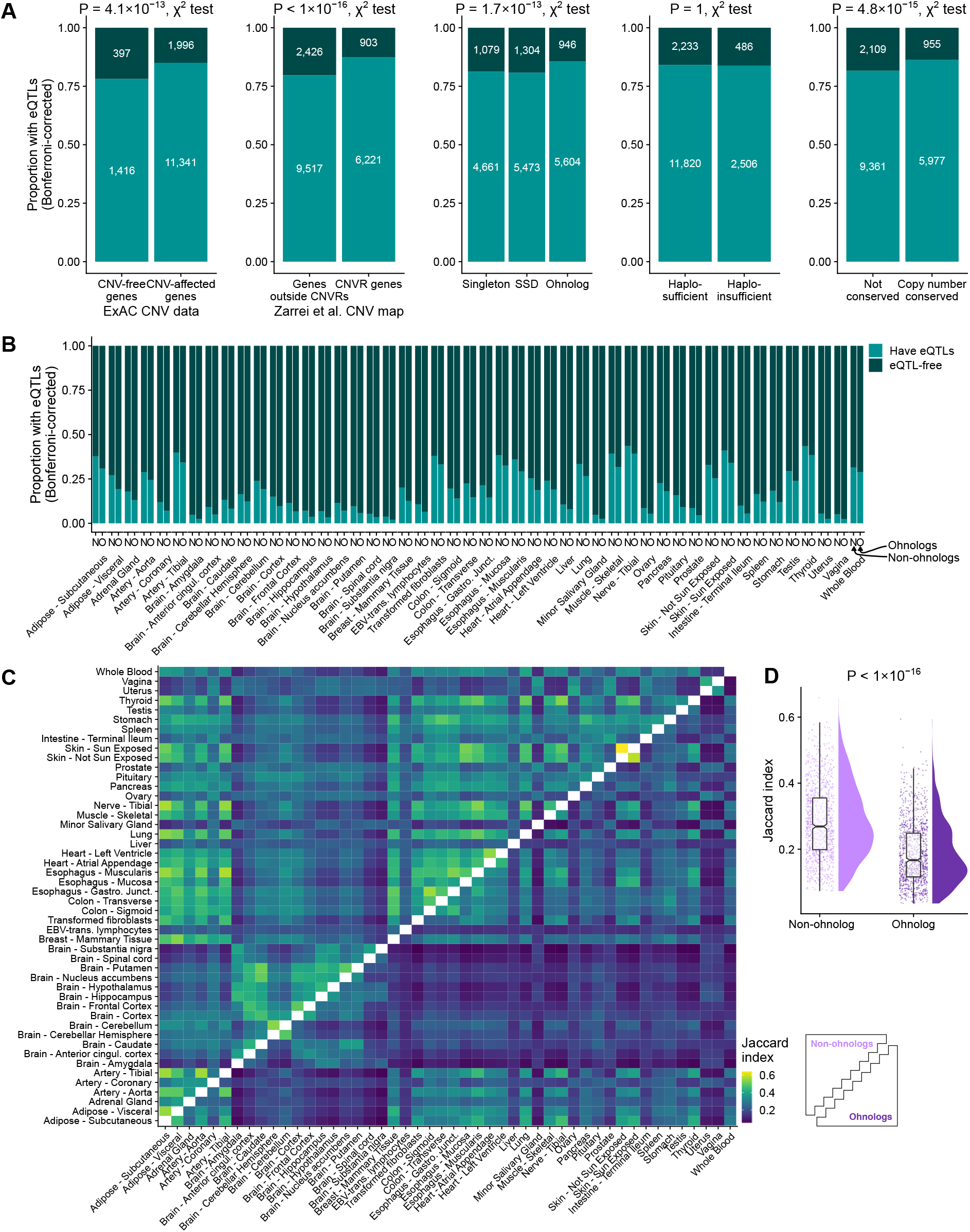
eQTL enrichment of CNVR genes and dosage sensitive genes. **A**, Proportion of genes affected by eQTLs for two sets of CNVs (ExAC CNV data and Zarrei et al. CNV map), ohnologs, haploinsufficient genes and mammalian copy number conserved (CCN) genes. P-values shown above each plot are Bonferroni-adjusted. **B**, Proportion of ohnologs (O) and non-ohnologs (N) affected by eQTLs per tissue. Sample size from 5,188-12,104. **C**, Pairwise overlap as Jaccard index between eQTL-affected genes in individual tissues. Upper triangle: Pairwise overlap of non-ohnologs; Lower triangle: Pairwise overlap of ohnologs. **D**, Distributions of pairwise Jaccard index for eQTL-affected genes between tissues for ohnologs and non-ohnologs.

While this first result suggests a straightforward correlation between lack of copy number constraints and presence of eQTLs, we found a contrary result with respect to long-term evolutionary copy number constraints (Figure 1A and Table 1). Ohnologs, which are generally refractory to further duplication and to CNV (Makino and McLysaght 2010) are enriched for being affected by eQTLs relative to non-ohnologs. Similarly, conserved copy number (CCN) genes, defined as genes which are in a one-to-one orthology relationships in 13 mammalian genomes (i.e. no gene loss or duplication within the mammalian tree), have also been seen to be refractory to CNVs (Rice and McLysaght 2017) and here are enriched for being affected by eQTLs.

The apparent contradiction between the dosage constraints operating on ohnologs across evolutionary timescales, and the enrichment for eQTLs demands further explanation. We considered the possibility that this might reflect something of the nature of the dosage constraints, specifically, whether or not it applied to all expressed tissues. Although ohnologs are more affected by eQTLs than non-ohnologs when considering all tissues together, within individual tissues we observe that, for every tissue tested, ohnologs are less frequently affected by eQTLs (Figure 1B). Given that the trend per tissue is the opposite to the trend observed when pooling tissues, we examined the possibility that more distinct subsets of ohnologs are affected by eQTLs in different tissues compared to eQTL-affected non-ohnologs (Figure 2).

**Figure 2.**
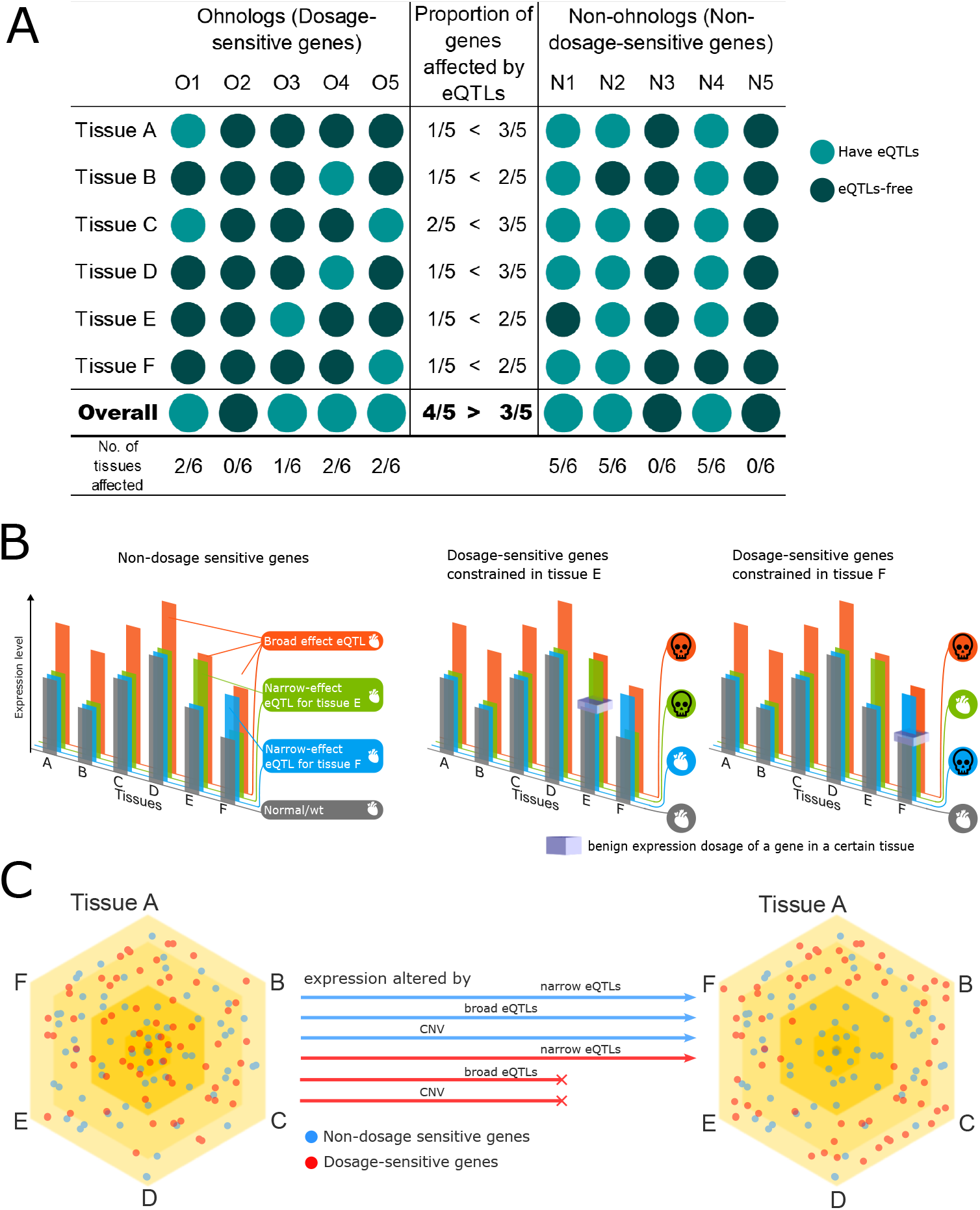
Differential predicted consequences of broad-effect and narrow-effect eQTLs on dosage-sensitive and non-dosage-sensitive genes across tissues. **A** A schematic representation of the proportion of genes affected by eQTLs globally and across individual tissues. In this hypothetical scenario, the ohnologs are more likely to be affected by an eQTL over all (4/5 compared to 3/5), but in each individual tissue they have fewer eQTLs. **B** Non-dosage sensitive genes tolerate expression alterations (left panel). Dosage constraints in some, but not all expressed tissues mean that broad effect eQTLS may be deleterious in dosage-sensitive genes, while narrow-effect eQTLs may or may not be tolerated, depending on the affected tissues (middle and right panels). Heart and skull icons are from Microsoft and are copyright and royalty free https://support.microsoft.com/en-us/office/insert-icons-in-microsoft-365-e2459f17-3996-4795-996e-b9a13486fa79 **C** Dosage-sensitive genes may be associated with narrow-effect eQTLs. The tissue specificity of eQTLs is illustrated, with broader eQTLs (affecting multiple tissues) located near the center and narrow-effect eQTLs (affecting specific tissues) positioned towards the periphery. Purifying selection, as shown in Figure B, leads to an enrichment of dosage-sensitive genes with narrow-effect eQTLs, while depleting those with broad-effect eQTLs or CNVs

The Jaccard index is a measure of similarity between sets and is the size of the intersection divided by the size of the union of the sets. If eQTL-affected ohnologs are more distinct between tissues compared to eQTL-affected non-ohnologs then we expect a lower Jaccard index between sets of ohnologs (i.e. a smaller overlap in eQTL-affected genes). We calculated pairwise Jaccard indices for eQTL-affected ohnologs between the 48 tested tissues, and similarly for eQTL-affected non-ohnologs (Figure 1C). We find a significantly lower similarity among eQTL-affected ohnologs compared to eQTL-affected non-ohnologs (median Jaccard index of 1,128 tissue comparisons of eQTL-affected ohnologs: 0.17 vs. 0.27 for non-ohnologs; *P <* 2.2 *×* 10^−16^, Mann-Whitney U test; Figure 1D).

### Duplication status, not expression level, predicts eQTL status per tissue

As ohnologs are more highly expressed than SSDs (median expression for ohnologs: 8.9 TPM vs. 6.0 TPM for SSDs; *P <* 2.2 *×* 10^−16^, Mann-Whitney U test, median expression for singletons: 10.3 TPM) and that genes affected by eQTLs tend to be more highly expressed (median expression in a tissue for eQTL-affected genes: 8.9 TPM vs. 7.9 TPM for unaffected; *P <* 2.2 *×* 10^−16^, Mann-Whitney U test), it was necessary to control for expression level when comparing ohnologs and nonohnologs for eQTL-enrichment. We binned genes into ten groups of equal size by their median tissue expression level across GTEx samples for each tissue. We observe that ohnologs are less frequently affected by eQTLs in every expression level category compared to non-ohnologs (Supp Figure **??**).

To investigate the contribution of a gene’s expression level and duplication status (ohnolog, SSD, singleton) to the presence or absence of an eQTL affecting a gene in a given tissue, we performed a logistic regression analysis. For each gene in a tissue, to predict its eQTL status, we used the gene’s median expression across GTEx samples in a tissue, and whether it is classed as an ohnolog, SSD, or singleton. We also included the interaction between expression level and duplication status in the model (Table **??**). From this logisitic regression analysis, it is clear that duplication status contributes far more to whether a gene is affected by an eQTL in a tissue than expression level. The odds of being affected by an eQTL for SSDs is 1.38 times that of ohnologs (*P <* 2.2 *×* 10^−16^), and for singletons is 1.41 times that of ohnologs (*P <* 2.2 *×* 10^−16^). Expression level and its interaction with duplication status, while each significant in the model, have odds ratios of 0.9998 and 1.0001 respectively and so meaningfully contribute little to eQTL status (*P* = 0.0003 for both).

### Dosage-sensitive genes have a smaller proportion of tissues affected by eQTLs

By definition, dosage-sensitive genes are under some form of dosage constraint in at least one of the tissues where they are expressed. CNVs may alter the amount of gene product across all tissues, which can be permissible in cases where the expression change is compatible with the constraint (e.g. a copy number gain of a gene that is haploinsufficient). However, an incompatible CNV in conflict with an expression constraint can produce a deleterious phenotype and will then be subject to purifying selection. eQTLs, on the other hand, can influence the expression of genes across a broad range of tissues or within only a single tissue and may thus avoid tissue-specific dosage constraints (Figure 2).

So far we have observed that ohnologs are enriched for being affected by eQTLs when considering all tissues simultaneously; are depleted for being affected by eQTLs when considering tissues individually; and that the tissues affected by eQTLs are more distinct between ohnologs than between non-ohnologs. Therefore, it follows that dosage-sensitive genes should have fewer eQTL-affected tissues per gene, presumably due to their levels being constrained in one or more of their tissues.

Examining this, we find that when comparing eQTL-affected genes, in each category of presumed non-dosage-sensitive genes we observe a higher proportion of expressed tissues affected by eQTLs than in the dosage-sensitive gene sets (Figure 3; Figure **??**; Table **??**).

**Figure 3.**
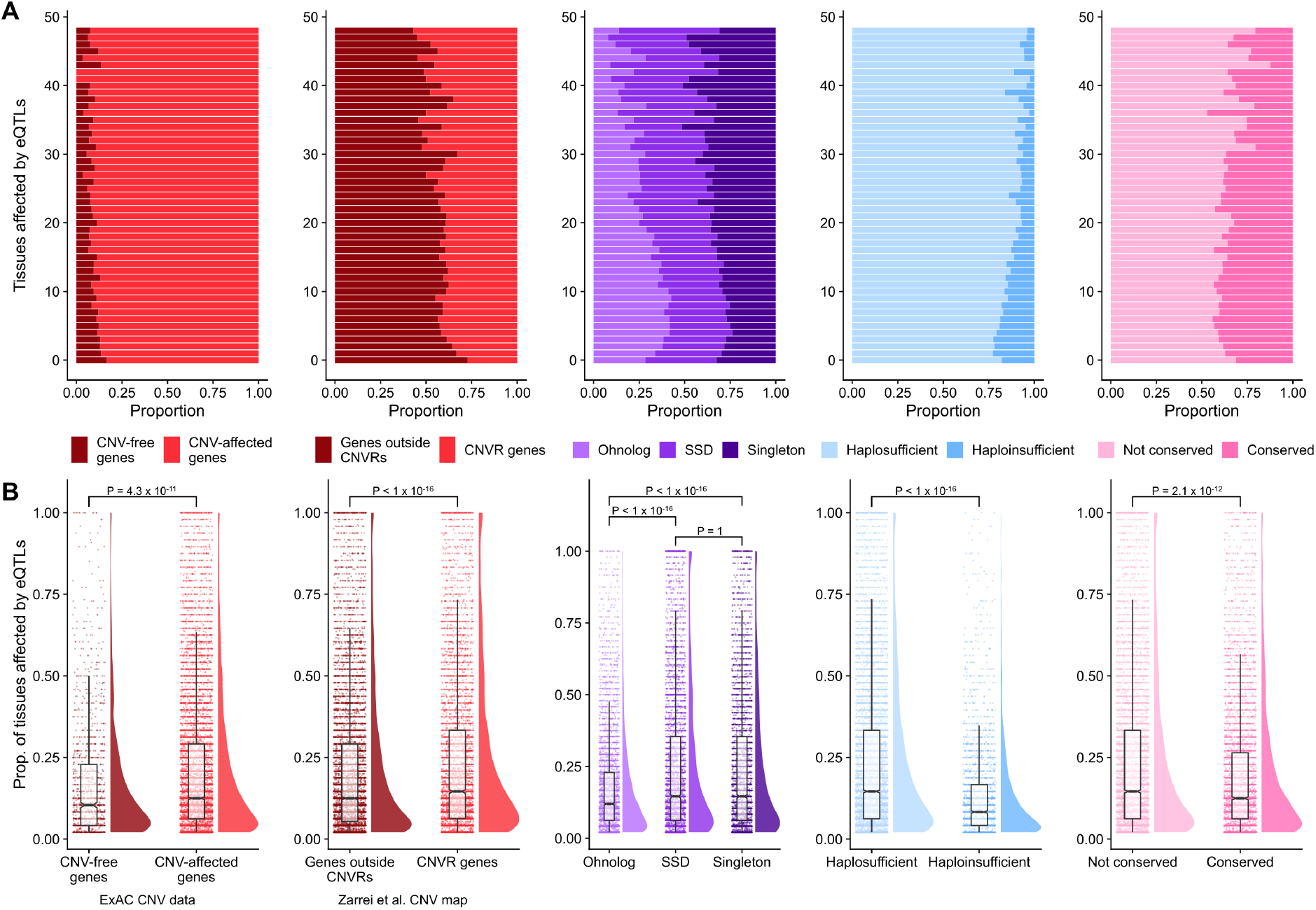
eQTL tissue specificity of dosage-sensitive genes. **A**, proportion of genes per number of tissues affected by Bonferroni-corrected eQTLs for genes affected by CNVs (red plots), ohnologs (purple), haploinsufficient genes (blue) and copy number conserved genes (pink). **B**, For each gene, proportion of tissues where the gene is expressed that are affected by Bonferroni-corrected eQTLs. P-values above each group for Mann-Whitney U tests and are Bonferroni-corrected.

### Dosage-sensitive genes are depleted for broad-tissue breadth eQTL

It makes intuitive sense that eQTLs that affect only a small number of tissues –narrow tissue breadth eQTLs– are less likely to clash with the dosage constraints of a given gene. To explore the relationship of eQTL tissue breadth and gene dosage constraints we focus on genes that are affected by (Bonferroni-corrected) eQTLs in at least 14 tissues. These genes could be affected by, say, 14 single-tissue eQTLs or one eQTL that affects expression in 14 tissues. This threshold was chosen as the top 10% of Bonferroni-corrected eQTLs affect gene expression in 14 or more tissues. We hereafter refer to these eQTLs affecting at least 14 tissues as broad-tissue breadth eQTLs. We then ask if dosage-sensitive genes within this set are depleted for being affected by broad-tissue-breadth eQTLs, even though they have a large number of eQTL-affected tissues.

We find no significant difference in the proportion of genes affected by broad tissue breadth eQTLs between genes experiencing CNVs and CNV-free genes (Figure 4). We do, however, observe that ohnologs are depleted for being affected by broad tissue breadth eQTLs compared to SSDs and singletons (63.4% of ohnologs vs. 74.9% for SSDs and 73.9% for singletons; *P* = 7.2 *×* 10^−9^, *χ*^2^ test). Haploinsufficient genes are not significantly different compared to haplosufficient genes for broad tissue breadth Bonferroni-corrected eQTLs and copy number conserved genes are significantly different from others after Bonferroni correction for multiple tests (67.9% of copy number conserved genes vs. 73.0% for genes not conserved; *P* = 0.02, *χ*^2^ test).

**Figure 4.**
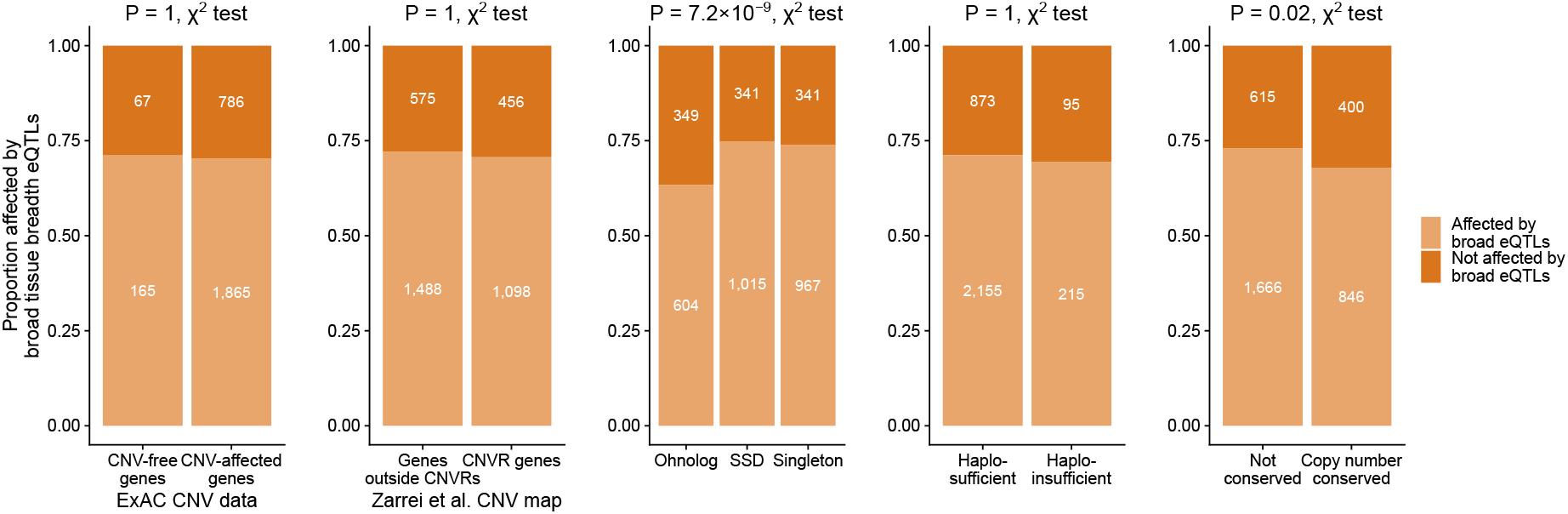
Broad tissue breadth eQTLs. Proportion of genes affected by broad tissue breadth Bonferroni-corrected eQTLs (influencing expression in 14 or more tissues) for two sets of CNVs (ExAC CNV data and Zarrei et al. CNV map), ohnologs, haploinsufficient genes and mammalian copy number conserved genes. *χ*^2^ test P-values shown above each plot are Bonferroni-adjusted.

In the Metasoft eQTL dataset the top 10% of eQTLs affect gene expression in 43 or more tissues, so we use this to define broad effect eQTLs to match the protocol for the first set. For these broad tissue breadth Metasoft eQTLs, CNV genes are not significantly different from CNV-free genes for both CNV datasets. However, ohnologs, haploinsufficient genes, and copy number conserved genes are all significantly depleted for broad tissue breadth Metasoft eQTLs (Figure **??**).

### eQTLs affecting dosage-sensitive genes have smaller effect sizes

The amount of influence an eQTL has on a gene’s expression level varies; some eQTLs only moderately increase or decrease mRNA level, while others have large effects. The direction and size of eQTL effects are quantified by the slope of the linear regression model used in identifying eQTLs in the GTEx project and represent the effect of the alternative allele relative to the reference allele. We hypothesise that dosage-sensitive genes may tolerate an eQTL of small effect while being refractory to eQTLs inducing larger expression changes.

To test this we compare the absolute value of the slope of eQTLs between our gene groups (Figure 5). We observe that CNV-free genes (median effect size: 0.35) and genes outside CNVRs (median: 0.36) both are affected by eQTLs with smaller effect sizes compared to CNV-affected genes (median: 0.38) and CNVR genes (0.45). Ohnologs, haploinsufficent genes and copy number conserved genes are all affected by eQTLs with significantly smaller effect sizes compared to their respective non-dosage-sensitive counterparts (Figure 5). As a more conservative test, rather than all eQTLs (22,715,646 eQTLs), we compare only the most significant eQTL for each gene per tissue (210,472 eQTLs; Figure **??**). We find the same significant trends in this more conservative set of eQTLs. We also compare allele frequencies from The 1000 Genome Project of SNPs associated with the most significant eQTL for each gene per tissue and find eQTLs affecting SSDs have a significantly higher allele frequency compared to eQTLs affecting ohnologs and singletons (Figure **??**). eQTLs affecting haploinsufficient genes and CNV-free genes both have significantly lower allele frequency than their counterparts.

**Figure 5.**
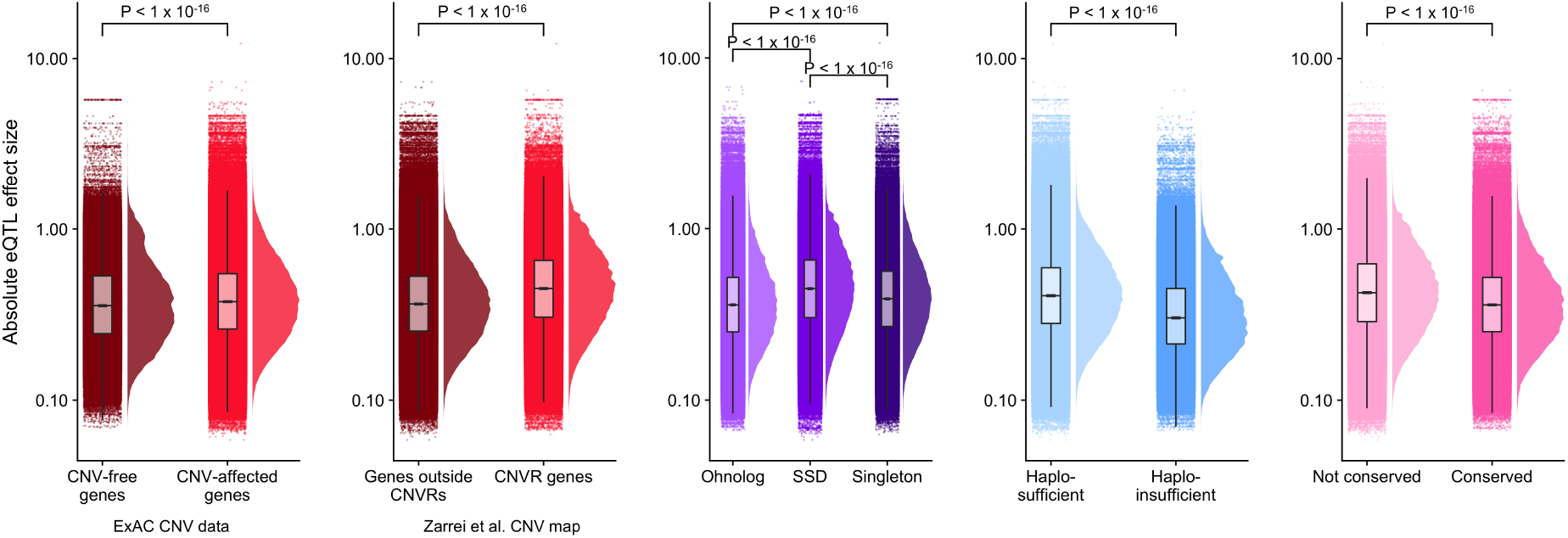
Absolute eQTL effect sizes for all eQTLs in different gene groups. Note the log10 scale. P-values above each group are for Mann-Whitney U tests and are Bonferroni-corrected.

## Discussion

The results presented here add a new dimension of complexity to our understanding of the consequences of dosage constraints on a gene’s evolution. Previous work has revealed an interesting and informative link between evolutionary gene duplicability and dosage sensitivity. Here we show that whereas ohnologs and copy-number conserved genes are less likely to be successfully duplicated over evolutionary times or within species, they are more likely to experience expression variation, as detected through eQTLs. At first glace, this would appear to contradict the interpretation of dosage sensitivity, however this can be explained as the difference between the system-wide and large increase caused by a gene duplication, compared to the possibility of localised and smaller-effect changes that can be achieved with eQTLs.

Using ohnologs, conserved-copy-number genes (CCNs) and genes without CNVs as proxies, we find that dosage sensitive genes, while generally more likely to be affected by eQTLS, are affected in a more tissue-specific manner, in proportionally fewer tissues, with smaller effects, and that SNPs linked to dosage sensitive genes are less likely to be eQTLs. We interpret this pattern of eQTL breadth and effect size as reflecting the dosage-sensitivity of the various classes of duplicate genes. Organism-wide or broad-effect eQTLs are likely to clash with the expression constraints of a dosage-sensitive gene, and ohnologs and mammalian copy-number conserved genes have previously been shown to be enriched for dosage-sensitivity.

One clear difference in these analyses is seen in the results obtained for evolutionary gene duplication status, and the results when considering CNVs. This may reflect two important differences between these types of duplication events. The first is that CNVs are often large enough to contain multiple genes, but the clinical effect of the CNV (benign versus pathogenic) may be driven by the presence of just one dosage sensitive gene in the region. This effect can create ‘CNV deserts’ in the genome, even if not all of the genes are in fact dosage sensitive (Makino, McLysaght, and Kawata 2013). This effect impacts these datasets because the CNV-free genes dataset will be a mix of dosage-sensitive genes and bystanders, and the dosage-sensitive genes may even be in a minority. We expect that this does not affect the evolutionary duplication status, where there has been sufficient time to resolve the dosage-constraints to a locus level with less linkage effect. Second, it is also known that CNVs can affect gene expression in complex ways (Franke et al. 2016) which may create extra layers of constraint and opportunity on this type of variation, and in ways which may not be entirely generalisable.

Taken together, our results suggest a complex interplay between the dosage constraints and the possible routes to variation in the amount of gene product. Whereas non-dosage-sensitive genes may vary in gene copy number and in gene expression level, due to their constraints this is not possible for dosage-sensitive genes, which can only vary in more restricted ways. Thus the only opportunities to vary the amount of protein produced from a dosage sensitive gene lie within tissue-restricted expression changes. This constraint channels the evolution of dosage sensitive genes towards this comparatively narrow evolutionary path. Detecting and interpreting these evolutionary patterns may shed new light on the functions and malfunctions of genes and the tissues where they are expressed.

## Methods

### Data

The data used in this paper’s analyses are obtained from publicly available data repositories.

All additional data are available at https://github.com/alanrice/paper-dosage-sensitivity-eqtl

#### Human eQTLs

Two datasets of eQTLs from The Genotype-Tissue Expression (GTEx) project V7 (GTEx Consortium 2017) were used: 1) significant single tissue SNP-gene associations for 48 tissues; 2) Metasoft eQTLs in 48 tissues. The first eQTL dataset of single tissue analyses was Bonferroni-corrected here for 48 tissues, and eQTLs were only further considered when they remained significant after correction in at least one tissue. The number of tissues where an eQTL affected expression was simply the count of tissues that remained significant after Bonferroni correction. The second eQTL dataset is derived from the first dataset of eQTLs where the data have been processed by Metasoft (Han and Eskin 2012) to give a posterior probability of being an eQTL in each of the 48 tissues. We included eQTLs when a tissue had a posterior probability of greater than 0.9. For this dataset, the number of tissues where an eQTL affected expression was considered to be the count of tissues with a posterior probability greater than 0.9.

#### CNV genes

Copy number variant regions were obtained from the inclusive CNV map in Zarrei et al. (2015) and a gene was considered to be intersecting with a region if any of the gene sequence was overlapped by one or more bases on either strand using Bedtools (Quinlan and Hall 2010). Genes that had a confident deletion or duplication call in 60,000 individuals from the Exome Aggregation Consortium (ExAC) release 0.3 dataset studied in Ruderfer et al. (2016) were defined as ‘CNV-affected genes’, otherwise genes were labelled ‘CNV-free genes’.

#### Whole genome and small scale duplicates, and singletons in the human and cow genomes

Singletons were defined as protein-coding genes that lacked a protein-coding paralog in Ensembl. A list of ohnologs (duplicates retained from whole genome duplication events early in the vertebrate lineage) were obtained from Singh and Isambert (2020) for both human and cow. Small scale duplicates were defined as protein-coding genes that had paralogs in Ensembl that were not classed as ohnologs. Ensembl version 75 was used for the human genome and version 96 for the cow genome.

#### Haploinsufficient genes

Haploinsufficient genes were defined as genes with a probability of loss-of-function mutation intolerance (pLI) of greater than 0.9 from the Exome Aggregation Consortium (ExAC) (Lek et al. 2016). For the purposes of comparison, only genes with available data and with pLI *<* 0.9 are included as ‘haplosufficient’.

#### Copy number conserved genes

Mammalian copy number conserved (CCN) genes are genes with no copy number changes in 13 mammalian genomes (Rice and McLysaght 2017).

#### SNP allele frequency

Allele frequencies from The 1000 Genome Project were downloaded from NCBI dbSNP for single nucleotide variants that corresponded to the most significant eQTL per gene/tissue (1000 Genomes Project Consortium et al. 2015; Sherry et al. 2001).

### Statistical analysis & figures

Unless otherwise stated, statistical tests were undertaken using R (R Core Team 2018) and figure plots were generated using ggplot2 (Wickham 2016).

Pairwise Jaccard index was calculated between each tissue for eQTL-affected ohnologs and nonohnologs seperately using the GeneOverlap R package (Shen 2020).

## Code availability

Jupyter notebooks (Kluyver et al. 2016) of analysis are available at https://github.com/alanrice/paper-dosage-sensitivity-eqtl.

## Notes

https://github.com/alanrice/paper-dosage-sensitivity-eqtl

### Competing Interest Statement

The authors have declared no competing interest.

