## Supplementary information for "Evolution of dosage-sensitive genes by tissue-restricted expression changes"

**Table S1.** Logistic regression model

| | *Dependent variable: Affected by eQTL (Y/N)* | | |
|---|---|---|---|
| | Coefficient (95% CI) | Odds ratio (95% CI) | P-value |
| Expression level | −0.0001 (−0.0002, −0.0001) | 0.9999 (0.9998, 0.9999) | 0.0003 |
| Duplication status - singleton | 0.3449 (0.3305, 0.3594) | 1.4119 (1.3916, 1.4325) | $< 2.2 \times 10^{-16}$ |
| Duplication status - SSD | 0.3231 (0.3090, 0.3373) | 1.3815 (1.3621, 1.4011) | $< 2.2 \times 10^{-16}$ |
| Expression level:singleton | 0.0002 (0.0001, 0.0003) | 1.0002 (1.0001, 1.0003) | $4.45 \times 10^{-6}$ |
| Expression level:SSD | 0.0001 (0.0001, 0.0002) | 1.0001 (1.0001, 1.0002) | 0.0003 |
| Constant | −1.7029 (−1.7134, −1.6925) | 0.1822 (0.1803, 0.1841) | $< 2.2 \times 10^{-16}$ |
| Observations | 790,141 (14,834-18,589 genes in 48 tissues) | | |

### Dosage-sensitive genes are affected by fewer eQTLs

The maintext analyses mainly concern proportions of tissues and eQTLs, however the absolute number of eQTLs affecting a dosage-sensitive gene could also be reduced if purifying selection is acting to remove deleterious variants that conflict with expression constraints. A similar pattern may also arise if dosage-sensitive genes are in regions of reduced polymorphism and as a result are affected by fewer eQTLs. GTEx test SNPs

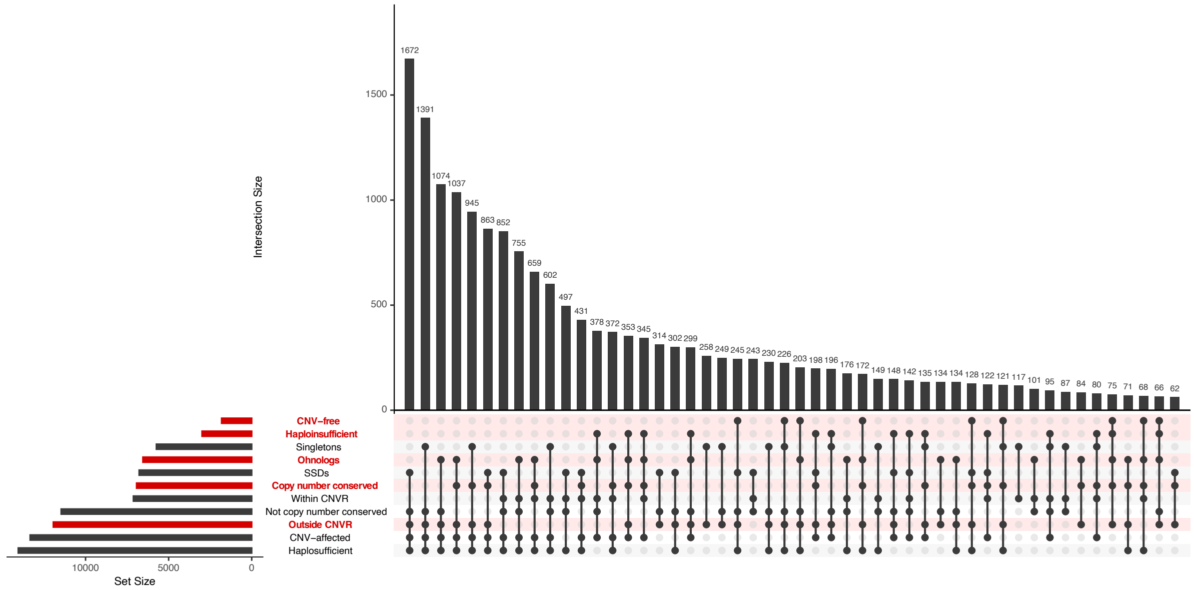


**Figure S1.** Counts of the overlaps between different gene sets used in these analyses. Gene sets that are broadly expected to be dosage sensitive are highlighted in red.

**Table S2.** Proportions of expressed tissues affected by eQTLs for eQTL-affected genes. P-values for Mann-Whitney U tests are Bonferroni-corrected for multiple tests.

| | | | Bonferroni-corrected eQTLs | | Metasoft eQTLs | |
|---|---|---|---|---|---|---|
| | | n | Median proportion affected tissues | P-value | Median proportion affected tissues | P-value |
| Zarrei et al. CNV map | Genes in CNVR | 7,124 | 14.6% | $< 1 \times 10^{-16}$ | 87.5% | $< 1 \times 10^{-16}$ |
| | Genes outside CNVRs | 11,943 | 12.5% | | 81.3% | |
| ExAC CNV genes | CNV-affected genes | 13,337 | 12.5% | $4.3 \times 10^{-11}$ | 85.1% | $< 1 \times 10^{-16}$ |
| | CNV-free genes | 1,813 | 10.4% | | 68.8% | |
| Duplication status | Ohnologs | 6,550 | 12.0% | $1 \times 10^{-16}$ | 72.9% | $1 \times 10^{-16}$ |
| | Small-scale duplications (SSDs) | 6,777 | 14.6% | | 87.5% | |
| | Singletons | 5,740 | 14.6% | | 91.7% | |
| Conserved copy number genes | Conserved genes | 6,932 | 12.5% | $2.1 \times 10^{-12}$ | 81.2% | $2.8 \times 10^{-8}$ |
| | Not conserved | 11,470 | 14.6% | | 85.4% | |
| Haploinsufficiency | Haploinsufficient genes | 2,992 | 8.3% | $< 1 \times 10^{-16}$ | 66.7% | $< 1 \times 10^{-16}$ |
| | Other genes | 14,053 | 14.6% | | 87.5% | |

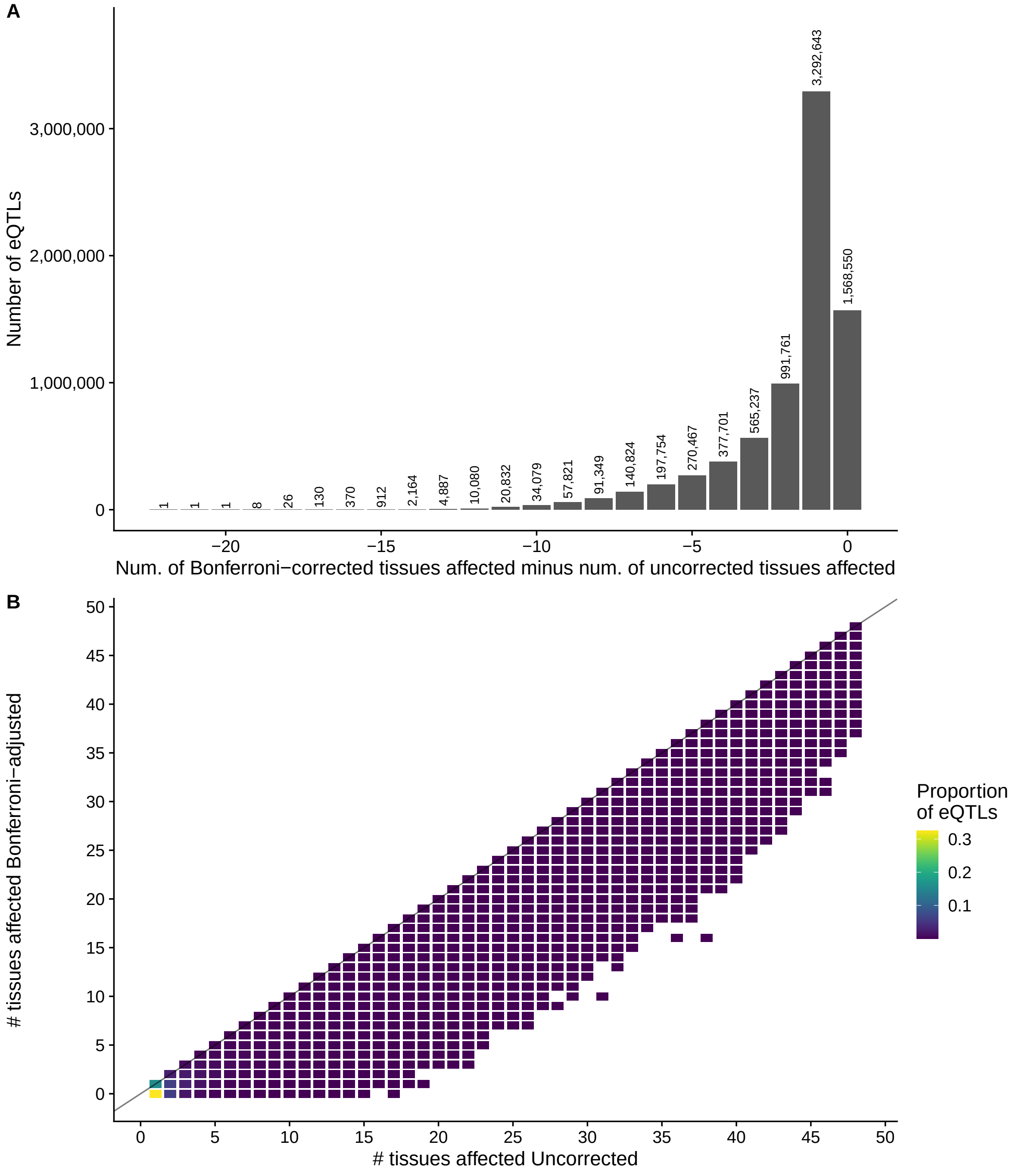


**Figure S2.** Number of tissues affected by each eQTL in GTEx v7 single tissue eQTL data (uncorrected) and after Bonferroni-correction for testing in multiple tissues.
**A** Difference in number of tissues affected between Bonferroni-corrected dataset and uncorrected dataset for each eQTL. **B** Heatmap of proportion of eQTLs with number of tissues in each dataset.

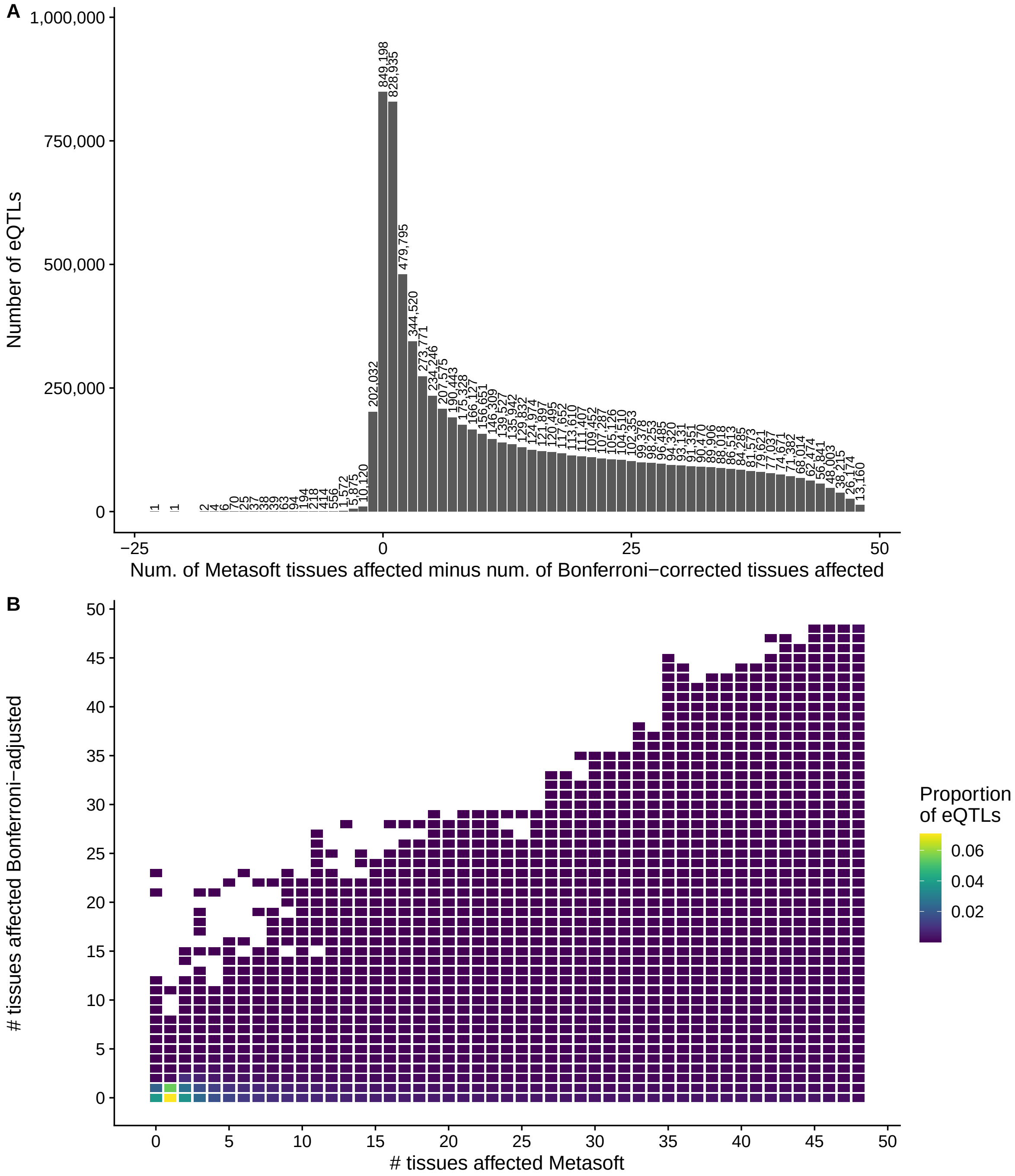


**Figure S3.** Number of tissues affected by each eQTL in GTEx v7 after Bonferroni-correction for testing in multiple tissues and GTEx v7 Metasoft eQTL dataset.
**A** Difference in number of tissues affected between Bonferroni-corrected dataset and Metasoft dataset for each eQTL. **B** Heatmap of proportion of eQTLs with number of tissues in each dataset.

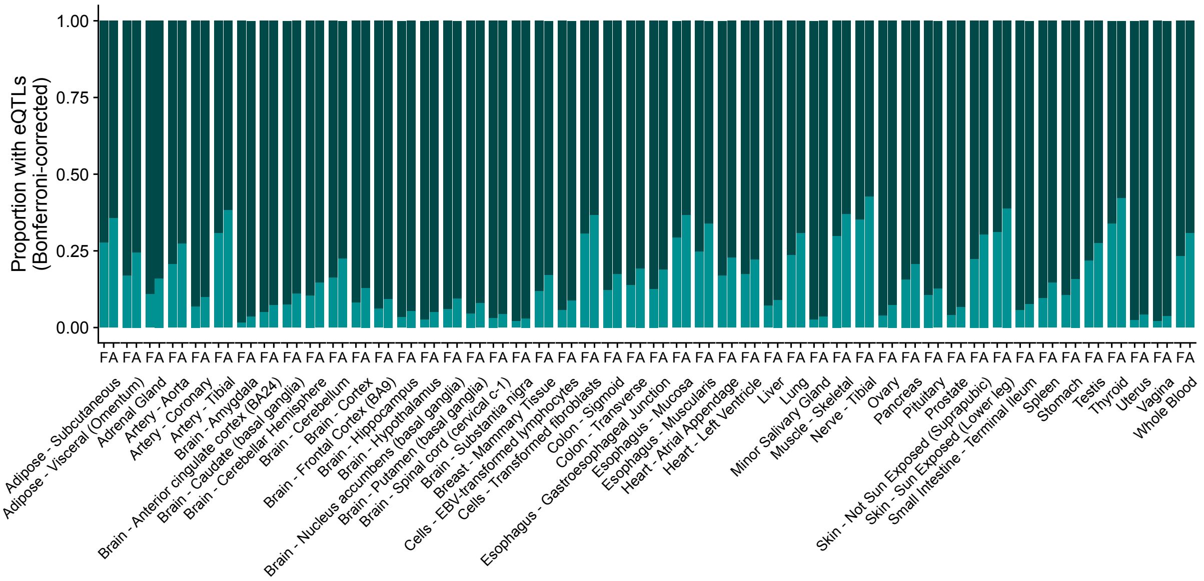


**Figure S4.** Proportion of CNV-free genes (F) have a lower proportion of eQTL-affected genes than CNV-affected genes (A) ffected by eQTLs per tissue.

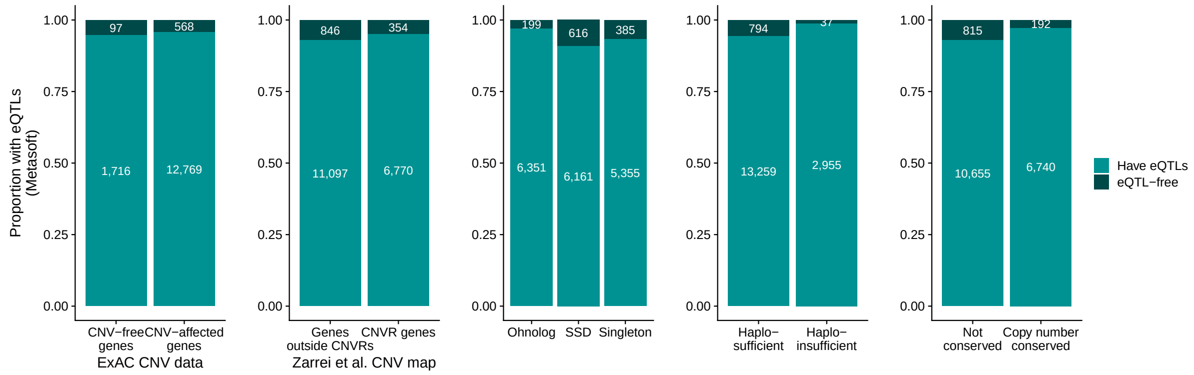


**Figure S5.** Proportion of genes affected by Metasoft eQTLs for two sets of CNVs (ExAC CNV data and Zarrei et al. CNV map), ohnologs, haploinsufficient genes and mammalian copy number conserved (CCN) genes.

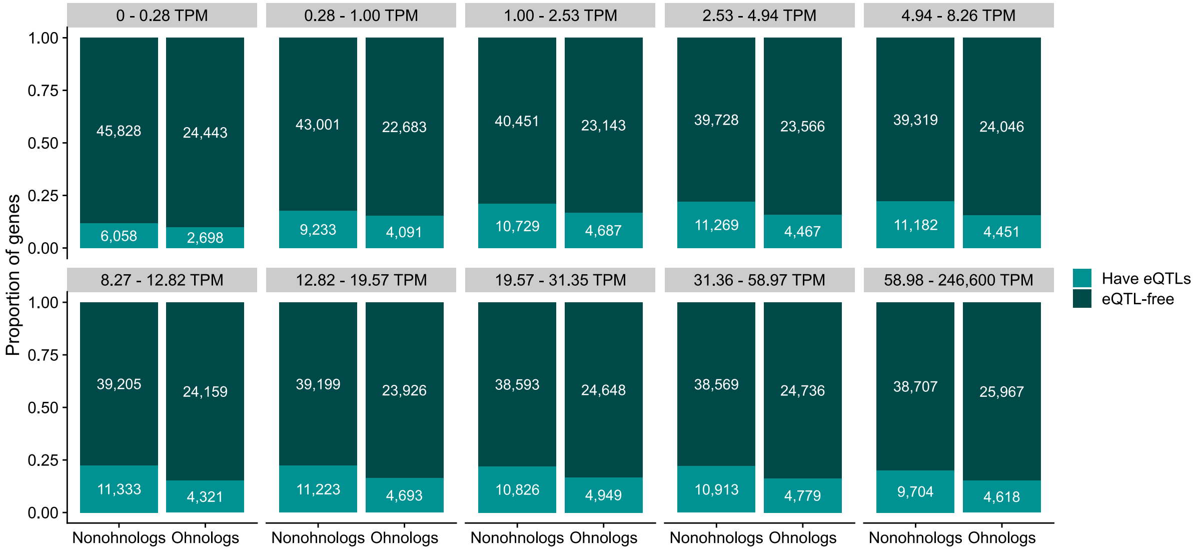


**Figure S6.** Proportion of genes affected by eQTLs grouped by median expression. The proportion of ohnologs and nonohnologs affected by significant Bonferroni-corrected eQTLs grouped by median expression per tissue into bins of roughly equal number. Above each bin, median expression values in transcripts per million (TPM).

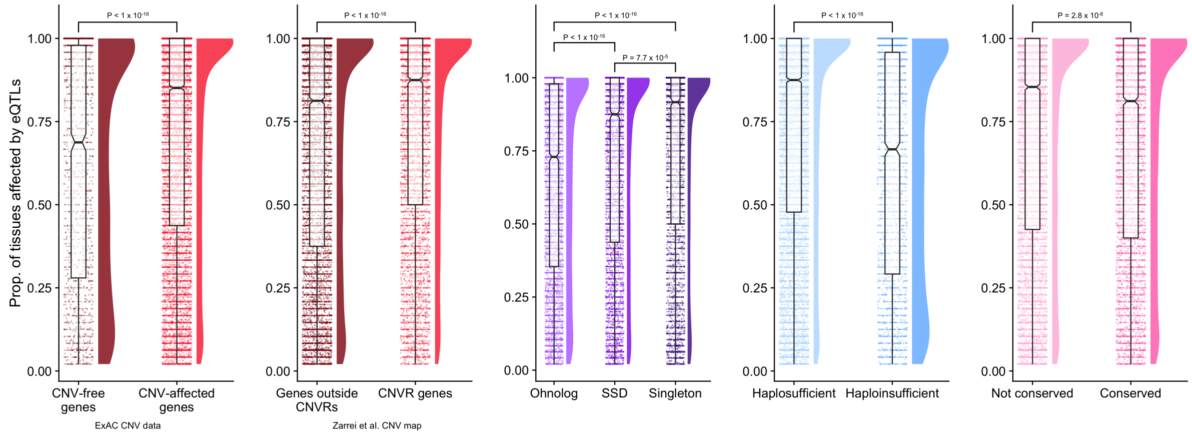


**Figure S7.** Proportion of expressed tissues that are affected by Metasoft eQTLs for eQTL-affected genes. P-values above each group are for Mann-Whitney U tests and are Bonferroni-corrected.

within 1 megabase (Mb) of the transcription start site of each gene for significant SNP-gene eQTL associations. The number of SNPs tested per gene range from 119 to 28,260. We observe a significant difference in the number of SNPs tested by GTEx between ohnologs and singletons (median SNPs ohnologs: 7,251; singletons: 7,371, respectively, $P < 1 \times 10^{-16}$, Mann-Whitney U test) and between SSDs and singletons (median SNPs SSDs: 7,236, $P < 1 \times 10^{-16}$) but no difference between ohnologs and SSDs ($P = 1$).

Due to the differing amount of polymorphism around genes, we compared the proportion of SNPs tested that are found to be significant eQTLs for our gene groups rather than the absolute number of eQTLs (Figure S8). As the number of eQTLs for a given gene varies between tissues, a gene can be included multiple times for every tissue where it has at least one significant eQTL. However, the number of tested SNPs within 1 Mb of a gene is constant between tissues.

We find that both ExAC CNV-affected genes and Zarrei et al. CNVR genes have a higher proportion of SNPs that are significant eQTLs (median: 0.0053 and 0.0054) compared to CNV-free genes and genes outside CNVRs (median: 0.0039 and 0.0051, $P < 1 \times 10^{-16}$ and $P = 6.8 \times 10^{-10}$, Mann-Whitney U test, respectively). Ohnologs have a lower proportion of SNPs that are significant eQTLs (median: 0.0042) compared to SSDs and singletons (median: 0.0055 and 0.0060, $P < 1 \times 10^{-16}$ for both, Mann-Whitney U test). SSDs and singletons are relatively similar ($P = 0.01$). Haploinsufficicent and conserved copy number genes also have a lower proportion of SNPs that are significant eQTLs compared to haplosufficient genes and genes that do not have conserved copy number (Figure S8). Similar trends are found for Metasoft eQTLs (Figure S9).

### Other mammalian genomes exhibit similar tissue-restricted expression changes

We wanted to explore whether the trends we observe for eQTLs affecting dosage-sensitive genes are unique to the human GTEx dataset and/or human genes, or are dosage-sensitive genes evolving by tissue-restricted expression changes observed elsewhere. To test this in

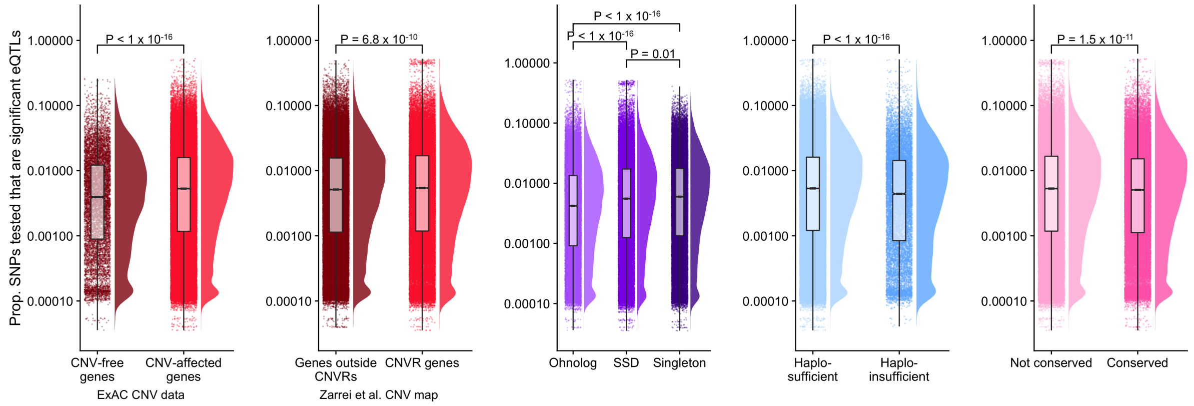


**Figure S8. Proportion of SNPs that are significant eQTLs.** Proportion of tested SNPs that are found to be significant Bonferroni-corrected eQTLs. As the number of eQTLs varies between tissues, each gene can be included multiple times, for every tissue where it has at least one significant eQTL. Note the log10 scale. P-values above each group are for Mann-Whitney U tests and are Bonferroni-corrected.

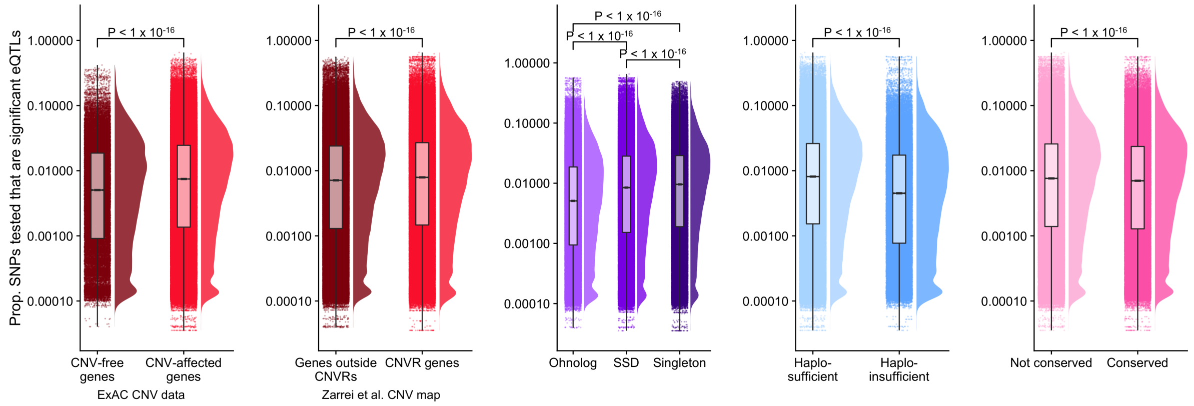


**Figure S9. Proportion of SNPs that are significant eQTLs.** Proportion of tested SNPs that are found to be significant Metasoft eQTLs. P-values above each group are for Mann-Whitney U tests and are Bonferroni-corrected.

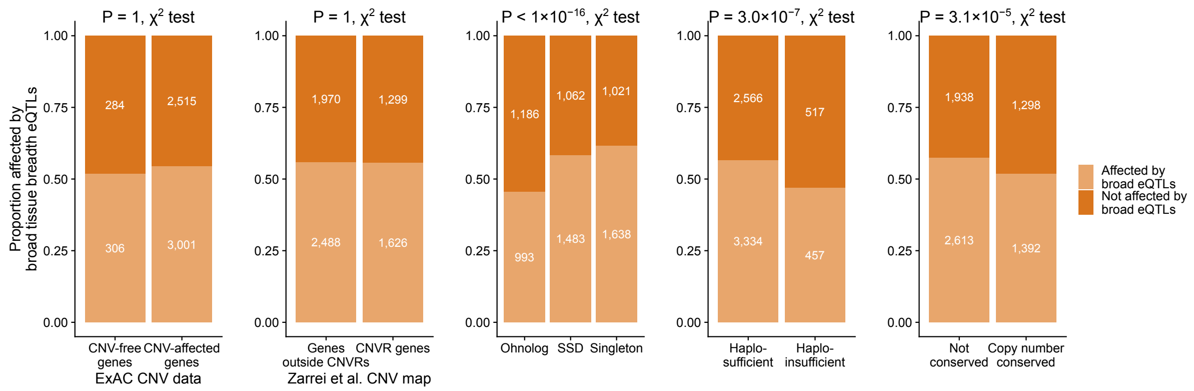


**Figure S10. Broad tissue breadth eQTLs** Proportion of genes affected by broad tissue breadth Metasoft eQTLs (influencing expression in 43 or more tissues) for two sets of CNVs (ExAC CNV data and Zarrei et al. CNV map), ohnologs, haploinsufficient genes and mammalian copy number conserved genes. $\chi^2$ test P-values shown above each plot are Bonferroni-adjusted.

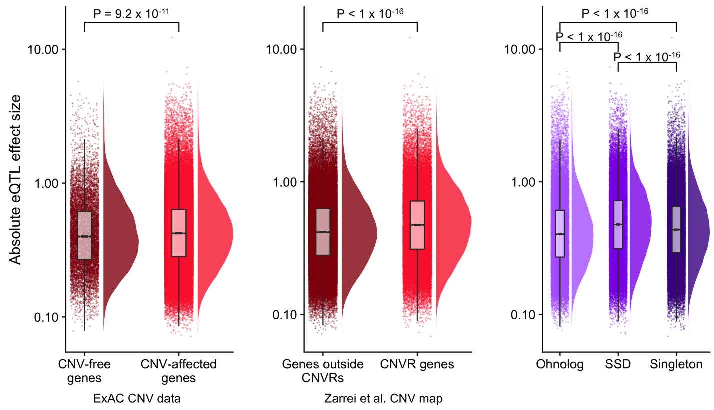


**Figure S11. Absolute eQTL effect sizes for most significant eQTL per gene/tissue in different gene groups.** Note the log10 scale and P-values above each group are for Mann-Whitney U tests and are Bonferroni-corrected.

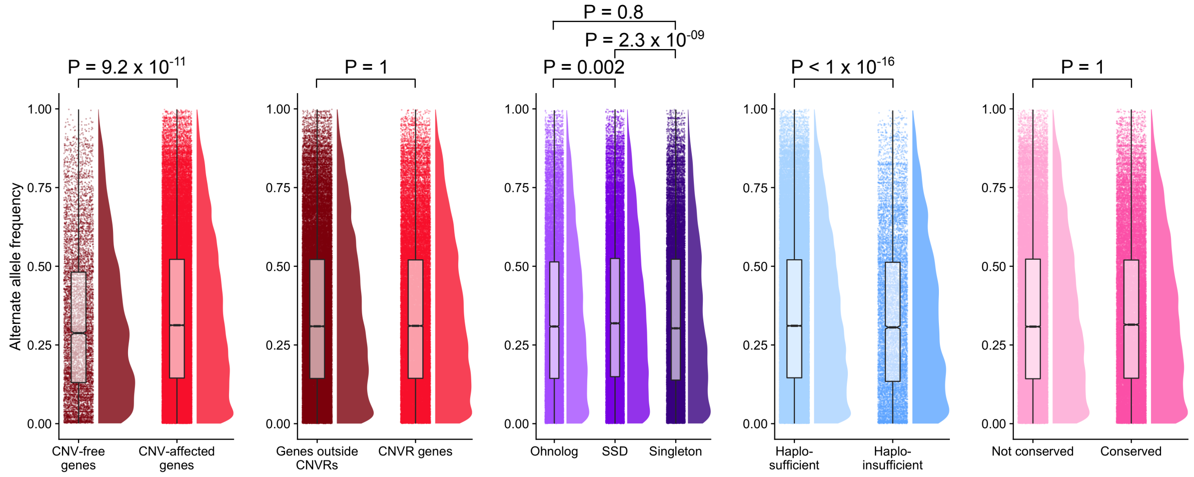


**Figure S12. Allele frequency of SNPs associated with most significant eQTL per gene/tissue in different gene groups.** P-values above each group are for Mann-Whitney U tests and are Bonferroni-corrected.

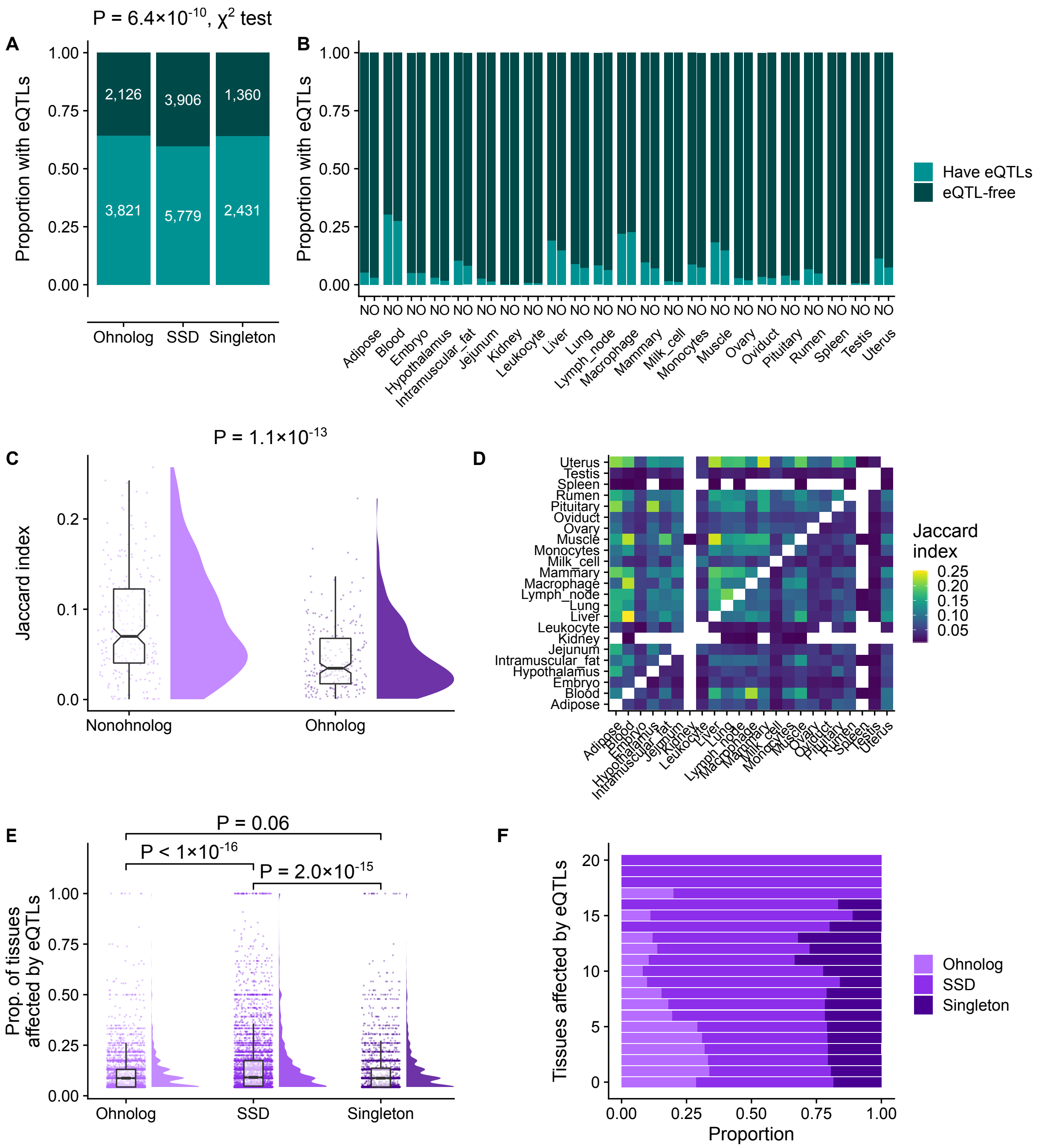


**Figure S13. eQTL trends for ohnologs and non-ohnologs in the cow genome.** **A** Proportion of cow ohnologs, SSDs, and singletons affected by eQTLs in any tissue. **B** Proportion of cow ohnologs (O) and non-ohnologs (N) affected by eQTLs per tissue. **C** Distributions of pairwise Jaccard index for eQTL-affected genes between tissues for ohnologs and nonohnologs. **D** Pairwise overlap as Jaccard index between eQTL-affected genes in individual tissues. Upper triangle: Pairwise overlap of non-ohnologs; Lower triangle: Pairwise overlap of ohnologs. **E** For each eQTL-affected gene, proportion of tissues where the gene is expressed that are affected by eQTLs. **F** Proportion of ohnologs, SSDs, and singletons per number of tissues affected by eQTLs.

another mammalian species, we employed a dataset of eQTLs from The Cattle Genotype-Tissue Expression atlas (cGTEx, Yao et al. 2022). This dataset was generated in a similar manner to the human GTEx project. Namely, cis-eQTL summary statistics were downloaded from The Cattle Genotype-Tissue Expression atlas (cGTEx) from `https://cgtex.roslin.ed.ac.uk/` (Yao et al. 2022). These were processed using the cGTEX script
4_cis-eQTL_p-value.nominal.correction_basePermutation.r to obtain a list of permutation corrected eQTLs that were subsequently Bonferroni-corrected for multiple testing for the number tissues tested.

We find that ohnologs in the cow genome are enriched for being affected by eQTLs when all tissues are considered together, similar to what we observe of human ohnologs (Figure S13A). We observe that 64.3% of cow ohnologs are affected by eQTLs in at least one of their expressed tissues compared to 59.7% of SSDs and 64.1% of singletons ($P = 6.4 \times 10^{-10}$, $\chi^2$ test). As observed in Figure **??**A, human ohnologs are enriched, and SSDs and singletons are depleted (standardised residuals of ohnologs: 7.9, SSDs: -4.8, singletons: -3.2), while in cow, singletons are also enriched (standardised residuals of ohnologs: 4.4, SSDs: -6.5, singletons: 3.1).

Considering individual tissues, ohnologs in the cow genome are less affected by eQTLs in 20 of 23 tissues/cell types (Figure S13B), with kidney, spleen and macrophage being the three tissues that differ from the trend. However, kidney and spleen have fewer than 20 eQTL-affected genes each so sample size might be a factor affecting the results. The remaining unusual tissue/cell type is macrophage (22.8% of ohnologs affected by eQTLs compared to 22.0% of non-ohnologs). When eQTL-affected genes are compared pairwise between tissues using the Jaccard index, eQTL-affected ohnologs are less shared between tissues compared to eQTL-affected non-ohnologs (Figure S13C and D), as was seen for the human data. We find significantly lower similarity among eQTL-affected ohnologs compared to eQTL-affected non-ohnologs (median Jaccard index of 220 tissue comparisons of eQTL-affected ohnologs: 0.04 vs. 0.07 for non-ohnologs; $P 1.1 \times 10^{-13}$, Mann-Whitney

U test).

As in the human dataset, we observe that ohnologs in the cow genome have a lower proportion of tissues affected by eQTLs compared to SSDs (median proportion of expressed tissues affected by eQTLs: 8.7% and 9.1%, respectively; $P < 1 \times 10^{-16}$, Mann-Whitney U test; Figure S13E and F). However, contrasting with the human data we observe that singletons in the cow genome have a lower proportion compared to SSDs but are not different from ohnologs (median proportion of expressed tissues affected by eQTLs: 8.7% and 9.1%, respectively; $P < 1 \times 10^{-16}$). In cow, singletons are not significantly different from ohnologs whereas in human, SSDs and singletons are similar and different from ohnologs.
